# Sexual signals of fish species mimic the spatial statistics of their habitat: evidence for processing bias in animal signal evolution

**DOI:** 10.1101/715177

**Authors:** Samuel. V. Hulse, Julien P. Renoult, Tamra C. Mendelson

## Abstract

The diversity of animal visual displays has intrigued scientists for centuries. Sexual selection theory has explained some of this diversity, yet most of this effort has focused on simple aspects of signal design, such as color. The evolution of complex patterns that characterize many sexual displays remains largely unexplained. The field of empirical aesthetics, a subdiscipline of cognitive psychology, has shown that humans are attracted to visual images that match the spatial statistics of natural scenes. We investigated whether applying this result to animals could help explain the diversification of complex sexual signaling patterns. We used Fourier analysis to compare the spatial statistics of body patterning in ten species of darters (*Etheostoma* spp.), a group of freshwater fishes with striking male visual displays, with those of their respective habitats. We found a significant correlation between the spatial statistics of darter patterns and those of their habitats for males, but not for females. Our results suggest that visual characteristics of natural environments can influence the evolution of complex patterns in sexual signals.

## Introduction

The diversity of visual patterning across animal species remains one of the most striking yet enigmatic of evolution’s puzzles. While visual patterns often function as camouflage, or evolve through other modes of natural selection, in many cases they are shaped by sexual selection. Although sexual selection is commonly invoked to explain the exaggeration of a sexual signal (e.g., Andersson 1994), little is known about why particular patterns are selected in some species, while different patterns are selected in others. This question becomes especially perplexing when closely related species exhibit a striking diversity of visual patterns, as in the peacock spiders of Australia, or the manakins of South America.

Some of the top candidate hypotheses explaining the evolution of signal design are the sensory bias and sensory drive models of sexual selection, which explain how environmental conditions can shape animal sensory systems, and thereby preferences for specific signal features^1–4^. These models have been especially useful for explaining the evolution of simple signal features, such as the color of a visual display, or the frequency spectrum of an auditory signal^1^. These features can be interpreted as components of signal efficacy, which refers to a signal’s ability to maximize information transmission^5^. In these models, the detectability of a signal determines its attractiveness, hence the central role of signal detection theory in sexual selection research. However, to date, little work addresses the question of how more complex traits, such as intricate visual patterns, can evolve through sensory drive^6^.

Recently, Renoult & Mendelson expanded the framework of sensory drive to include efficient information processing as an explanation for complex signal design^5^. Efficiency describes the transmission of information at low metabolic cost. The expanded sensory drive framework posits that the neural circuitry underlying sensory perception is tuned to efficiently process habitat-specific features, and that this specialization can lead to preferences for particular visual patterns, as might be displayed by a potential mate. This hypothesis is grounded in information theory^7^, specifically the efficient processing hypothesis of Horace Barlow^8^. Action potentials are metabolically expensive, and neural systems reduce metabolic expenditure by reducing the number of action potentials required for signal processing^9^. This reduction is accomplished by leveraging the statistical redundancies in sensory stimuli to create a “sparse” neurological representation (or code); that is, for any given stimulus, relatively few neurons are active at any given time, and those active neurons are highly tuned to the redundant (regular) features of the perceptual environment^10^. Visual information in particular contains a great deal of statistical redundancy, or regularity, that visual systems have adapted to efficiently process^11–13^. Similar adaptations for efficiency have also been shown in auditory and olfactory processing^14,15^.

The field of human empirical aesthetics uses Barlow’s efficient processing hypothesis to explain why humans find certain visual stimuli, like works of art, more appealing than others^16^. A number of studies have found that humans prefer, and find more pleasurable, images that are more efficiently processed^17–19^. In parallel, other studies have shown that visual art has fractal-like statistics similar to natural scenes, whereas less aesthetic images, such as those of laboratory objects (i.e. spectrometers, lab benches, etc...) do not. Psychologists hypothesize that people prefer art with the spatial statistics of natural scenes because our brains have evolved to efficiently process them^20–24^. Results from cognitive psychology therefore suggest a processing bias rooted in the reward (pleasure) of efficient information processing^5,25^. Renoult & Mendelson hypothesize that this processing bias is not limited to humans, predicting that other animals should also prefer the fractal-like statistics of their habitats, and that this preference could help explain the evolution and diversification of complex animal signal patterns^5^.

To date, psychological studies of efficient processing have considered natural scenes to be homogeneous, disregarding potential variation between habitats. However, other studies have shown that habitats can differ significantly in spatial statistics, and specifically in the statistics that measure visual redundancies^26,27^. Such studies quantify the intuitive: an image of the forest understory, with highly repeating vertical contrast (trees), will have different spatial statistics than an image of a desert, or a beach. Thus, organisms occupying habitats with different spatial redundancies are predicted to have environment-dependent differences in visual processing. In keeping with a hypothesis of sensory drive, these processing differences could lead to environment-dependent differences in pattern preferences, with the most efficiently processed (and thus preferred) environments being those inhabited by a given perceiver or by its ancestors. A hypothesis of signal diversification based on processing bias therefore predicts that the spatial statistics of complex visual signals whose function is attraction, as in courtship, will match those of the local habitat^5^. Here, we test that prediction in a diverse genus of freshwater fish with complex visual courtship signals.

Darters (Percidae: *Etheostoma*) are an especially appropriate system in which to study the diversity of visual patterns in the context of sensory drive and processing bias. Darters are a diverse group of benthic freshwater fish, found throughout the eastern United States^28^. Phylogenetic evidence suggests that the most recent common ancestor of darters existed between 30 and 40 million years ago, and darters are the second most species rich group of freshwater fish in North America^29,30^. During their breeding season (typically March through May), male darters of most species exhibit species-specific nuptial coloration used in courtship and competition, while females typically remain drab and cryptic. In addition to their striking male color displays, different species of darters exhibit marked variation in patterning (Figure 1). Mate choice assays in some darter species have shown that both males and females prefer the nuptial coloration and pattern of conspecifics^31–35^. While the most closely related darter species have similar habitat preferences, distantly related species have divergent habitat preferences that distinguish many sympatric species within a community^36–38^. These distinct habitats could exhibit distinguishable spatial statistics that might drive divergence in pattern preferences and ultimately divergence in the patterns of male sexual signals.

**Figure 1.**
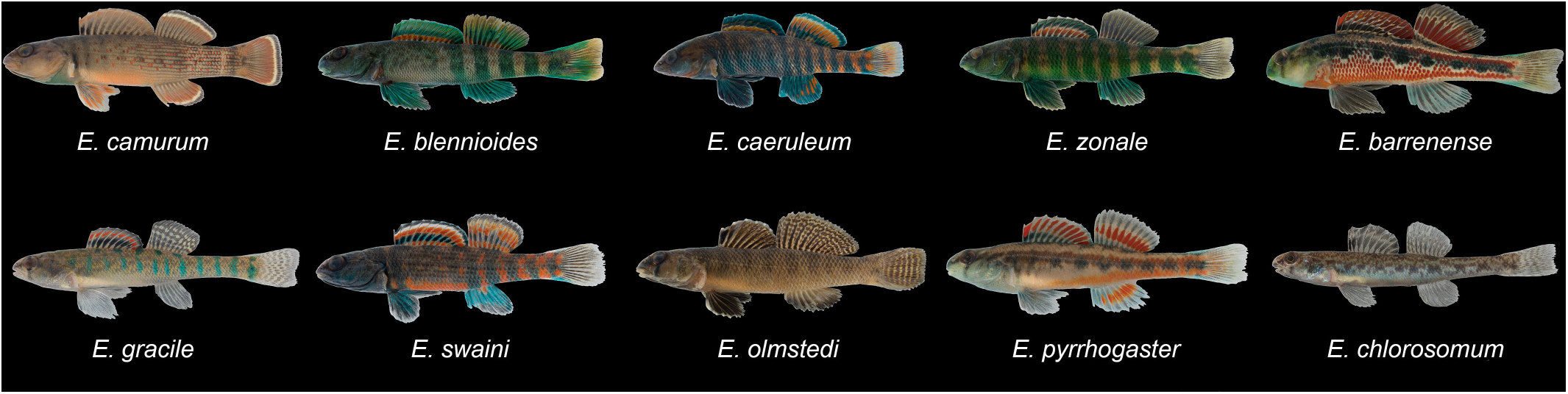
Example images for males of each species included in this study.

In this study, we investigated for the first time whether differences in environmental visual statistics correlate with observable differences in the visual statistics of sexually selected phenotypes. We captured digital images of ten species of darters that occur in five different classes of aquatic habitats, as well as images of their habitats. We used Fourier analysis to characterize the spatial statistics of habitats and the darter species that reside in them and tested for a correspondence between these statistics. Fourier analysis, which decomposes a signal into its component frequencies, is one of the most commonly used methods for analyzing visual images and a central tool in the field of empirical aesthetics^23,24,39–41^. For visual images, it indicates how luminance contrast (i.e., energy) is distributed across a range of spatial sinusoidal frequencies. Lower spatial frequencies correspond to large scale features in an image, such as the horizon line; higher spatial frequencies correspond to fine scale features, such as grains of sand. When plotted on log-log axes, the relationship between frequency and energy can be approximated by an affine function; the slope of this function is referred to as the slope of the Fourier power spectrum (1/*f;* hereafter, “Fourier slope”; Figure 2). Many studies use the Fourier slope to examine similarities between aesthetically pleasing images and natural scenes^18,21,42^. We applied these methods to darter color patterns and their preferred habitats, testing a hypothesis of sensory drive.

**Figure 2.**
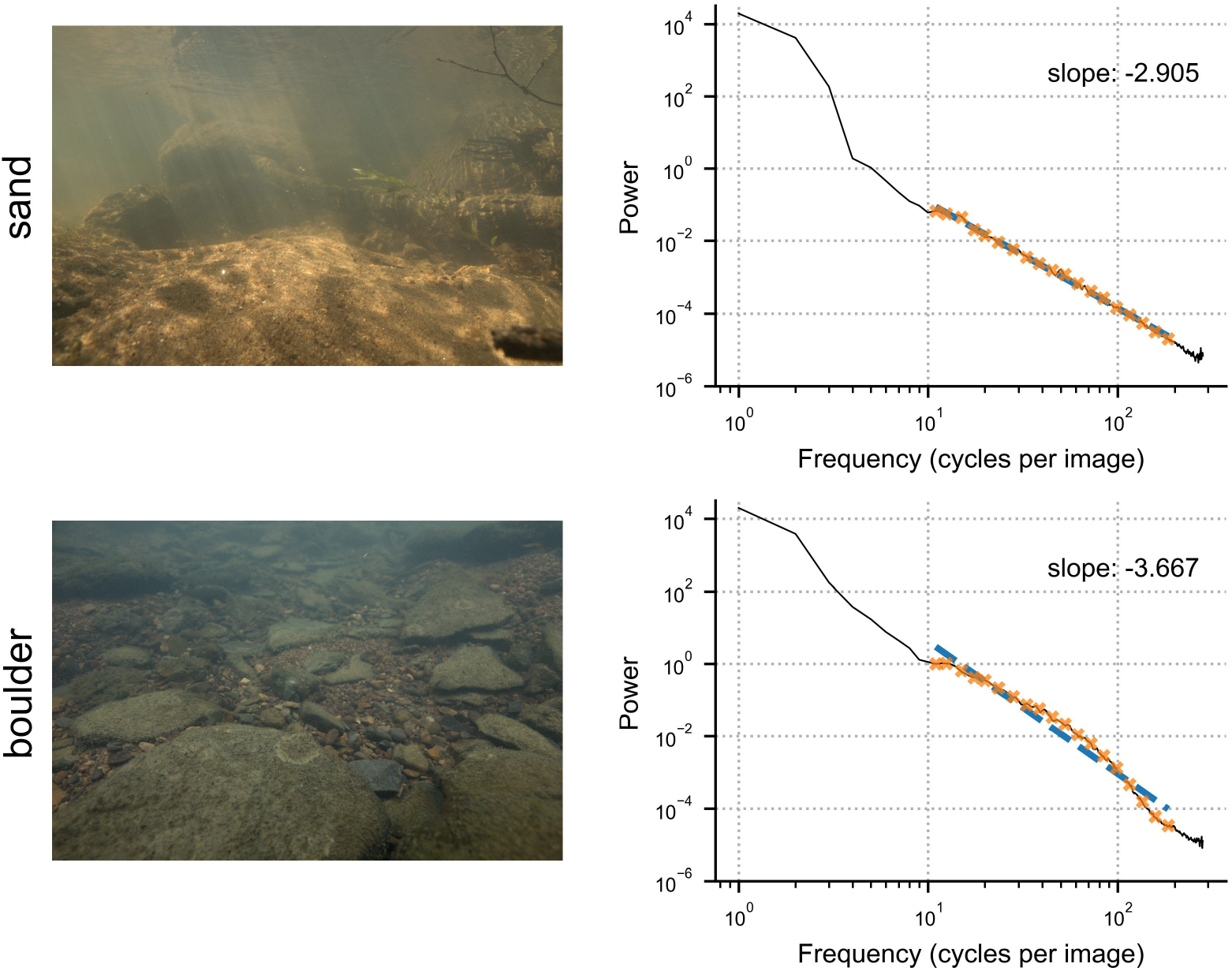
The Fourier slope of images from two different habitat types. The blue line represents the entire power spectrum, while the orange x’s are the bin locations. The black line is the fit of the bins, and represents the slope used for this analysis. For habitat images, the bins are evenly distributed between 10 and 200 cycles per image.

## Results

### Slope of the Fourier power spectrum

We found significant variation in the Fourier slope across species for both males (p = 2e-16, n = 302, F = 16.2, df = 9, Figure 3) and females (p = 3.47e-9, n = 274, F = 7.08, df = 9, Figure 3). Overall, males had a higher slope than females (males: −3.09 +/ −0.342 SD, females: −3.27 +/- 0.306 SD, n = 576); this difference was significant in 5 out of 10 species (two-tailed Student’s t-test with Bonferroni correction, species’ results provided in Supplementary Table 2). Differences in the Fourier slope across habitat classes were also significant (p = 2e-16, n = 2388, F = 58.8, df = 4, Figure 4).

**Figure 3.**
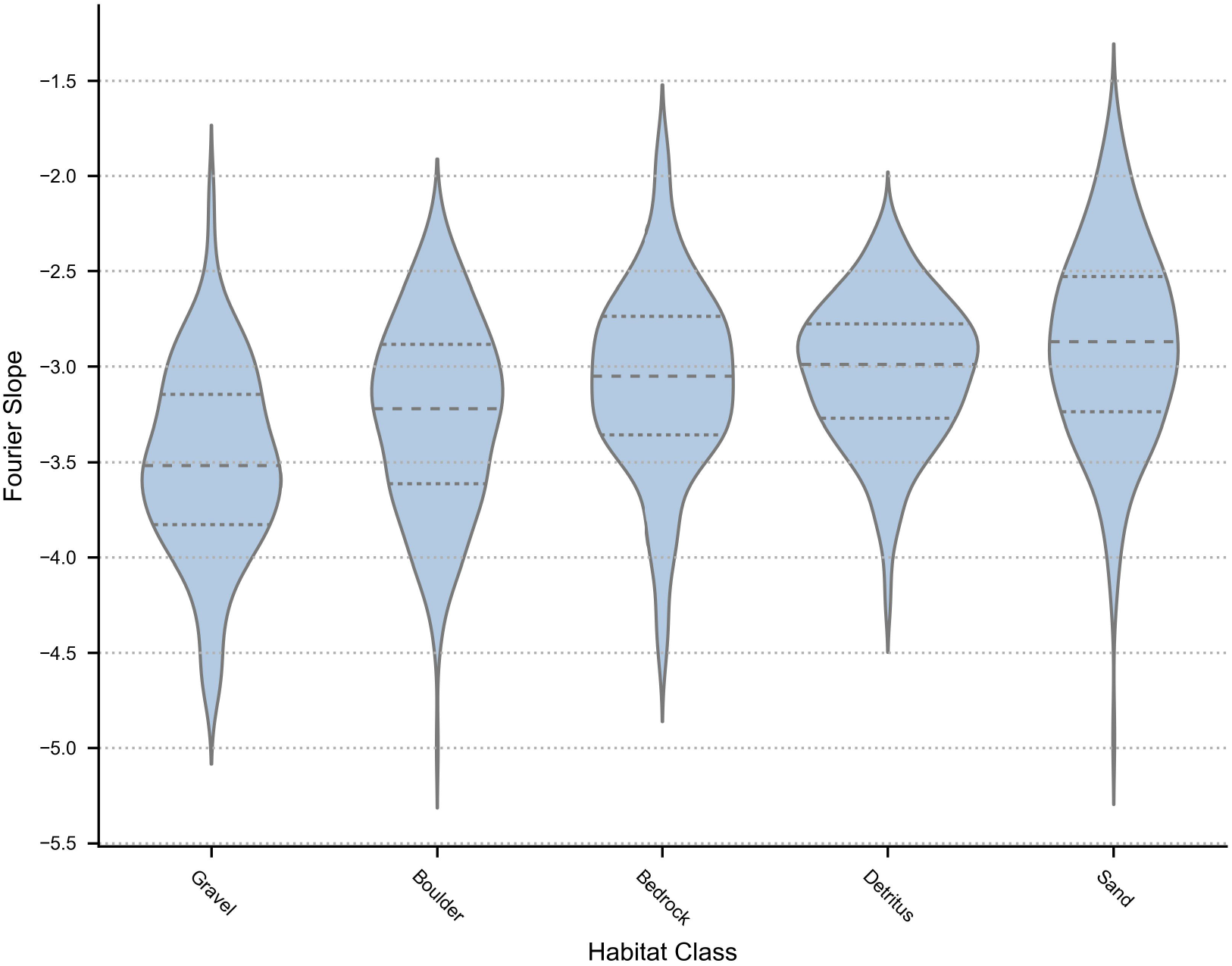
Distribution of Fourier slopes for males and females of ten species in the genus *Etheostoma*. Dashed lines represent the median for each group, and dotted lines represent the interquartile range. The left (blue) half of each violin plot are values for males; the right (orange) half of each violin plot are values for females. Species are grouped by habitat type.

**Figure 4.**
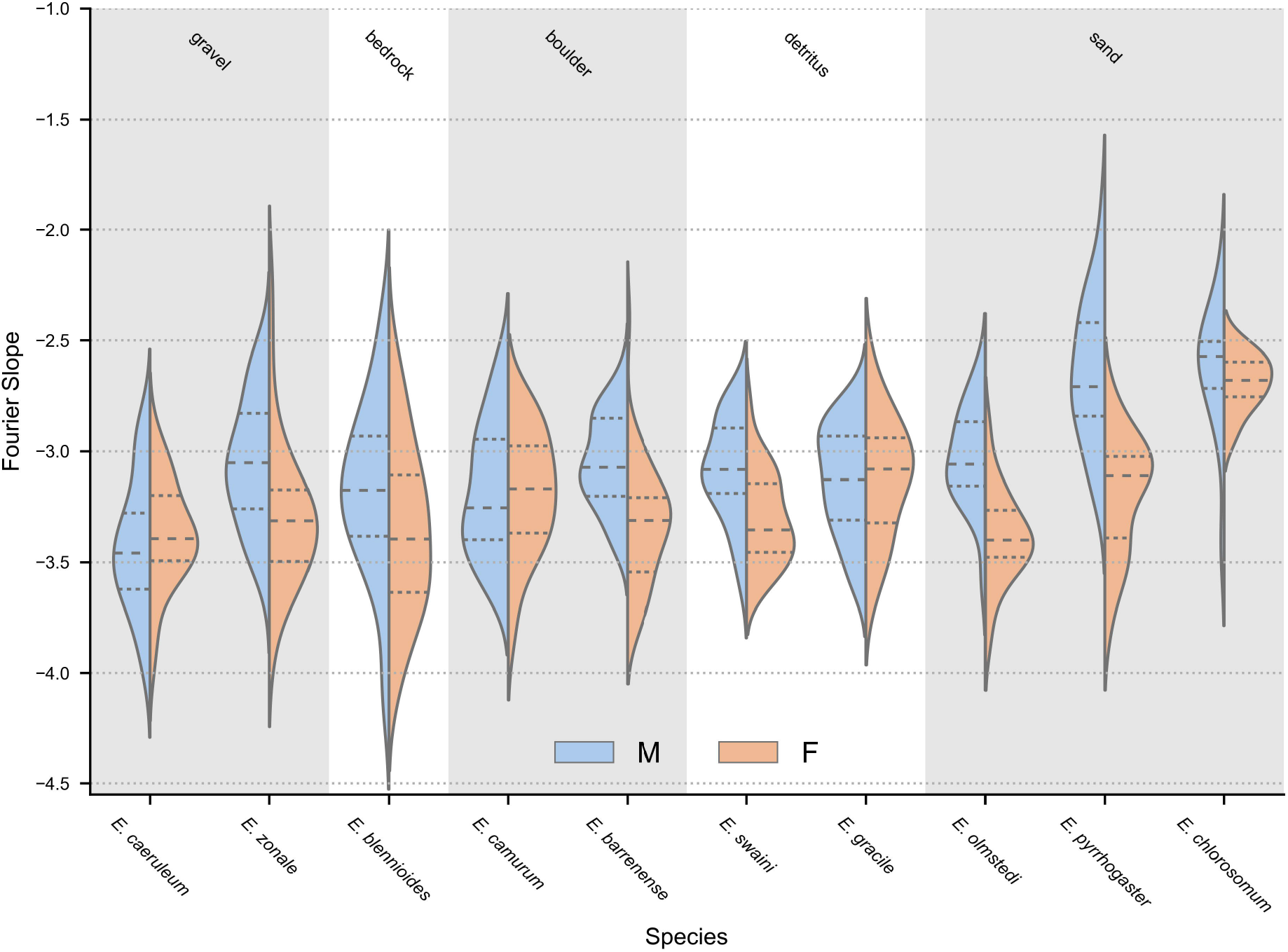
Distribution of Fourier slopes for images of five habitat types. Dashed lines represent the median for each group, and dotted lines represent the interquartile range.

Our model of darter Fourier slopes included habitat slope as a fixed effect, as well as phylogeny and capture site as random effects (models without both random effects had a higher DIC). For the following results, the mean effect size (β) is reported with its 95% credible interval (CI). Results where the CI includes zero indicate a statistically nonsignificant result with α = 0.05. The correlation between the Fourier slope of color patterns and that of their corresponding habitat classes was significant for males (β = 0.436, CI = [0.0442, 0.834], pMCMC = 0.0262, DIC = 60.4, Figure 5a), but not for females (β = 0.295, CI = [−0.389, 1.03], pMCMC = 0.3709, DIC = 36.72, Figure 5b). We did not find a strong effect of the sample site on the Fourier slope of the fish for either males or females (males: β = 0.0327, CI = [0.0111, 0.0594], females: β = 0.0323, CI = [0.0089, 0.0618]). The effect of phylogeny was minimal for males and its exclusion from our model did not change the significance of our results (β = 0.00516, CI = [1.5e-17, 0.0252]). Additionally, for females the effect of phylogeny was minimal (β = 0.0467, CI = [0.0023, 0.121]), and its exclusion from our model did not produce a significant correlation between the Fourier slope of females and that of their habitats.

**Figure 5.**
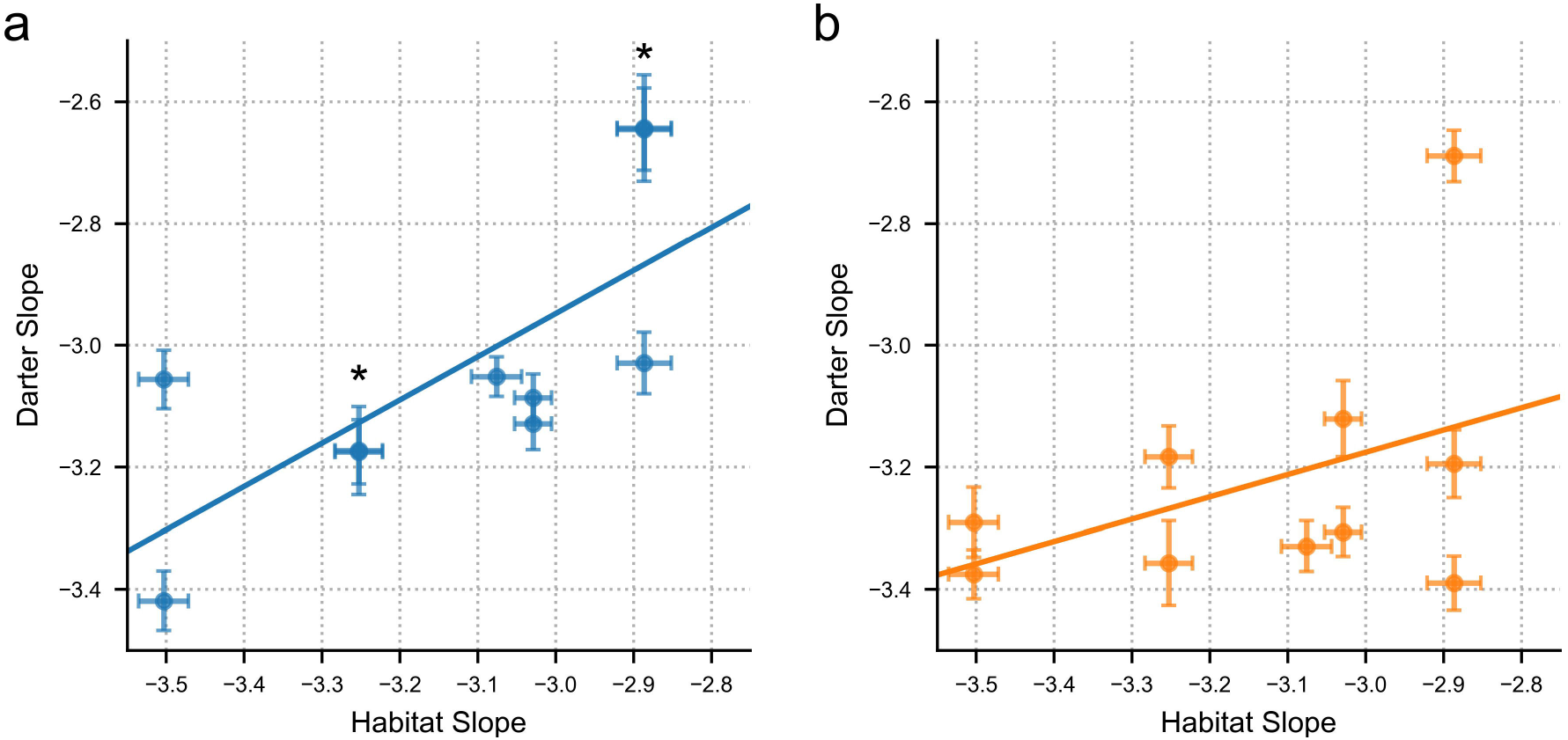
Scatterplots comparing mean Fourier slopes of fish versus habitat for males (a) and females (b) in ten species of *Etheostoma*. Error bars represent the standard error. For the males, *E. pyrrhogaster* and *E. chlorosomum* have nearly identical values, and appear superimposed on top of each other. This is also the case for *E. camurum* and *E. blennioides*. These species are marked by (*).

## Discussion

Sensory drive posits that animal signals are shaped by the environments in which they are transmitted^1,3,4^. Environmental features affect not only signal transmission but also the sensory and perceptual systems of receivers, which in turn can affect the course of signal evolution. Therefore, one of the central predictions of sensory drive is that signals evolving due to sexual selection will vary predictably with the environment in which they evolve^3^. Our results support this prediction. We have shown that the habitats occupied by different darter species have different visual statistics, measured as the distribution of luminance contrast across different spatial frequencies (i.e., Fourier slope). We also found a significant correlation between the Fourier slope of a species’ habitat and that of male nuptial patterns. Our results are therefore consistent with a hypothesis of sensory drive that incorporates efficient information processing in receivers as a driving force in preference and signal evolution^5^.

The framework established by sensory drive is crucial to our understanding of sexual selection, because it can explain not only why signals become elaborate, but also why they take their particular form^1,3,4,43^. To date, however, that framework has been rooted primarily in signal detection theory, which does not account for more complex features integrated at higher levels of perceptual processing. In a review of the state of animal coloration research, Endler and Mappes noted that the effect of perceptual processing on the evolution of color patterns is virtually unknown^6^. In addition, of the 154 studies investigating sensory drive analyzed by Cummings and Endler, none leveraged perceptual processing to understand how spatial structure in visual patterns evolve^1^. Instead, the bulk of research focused on signal detection via retinal photoreceptors. Information theory is a broader context that motivated the efficient processing hypothesis of Barlow^8^, which emphasizes energy efficiency in information processing. Rooting the framework of sensory drive in information theory thus suggests a novel way forward for understanding how complex sexual signals evolve and a new mechanism through which sensory drive can shape signal design.

Extending sensory drive to include processing efficiency is motivated by work in empirical aesthetics, which demonstrates that people prefer images that match the spatial statistics of natural (terrestrial) scenes. Much of this work is based on the Fourier slope. Images of terrestrial habitats tend to have a Fourier slope around −2 with surprising regularity. In contrast, images of non-natural items (e.g., anthropogenic landscapes and objects) have a Fourier slope that deviates from −2^44,45^. One notable exception is works of art, which tend to have a slope similar to that of natural terrestrial habitats ^20,23,42^. For instance, Redies et al. found that artists’ portraits of human faces mimic the Fourier slope of natural landscapes more closely than images of human faces do^21^. Beyond works of art, people also tend to prefer synthetic images with a Fourier slope around −2^40^.

A preference for Fourier slopes characteristic of natural habitats is thought to arise from the pleasure of efficient processing, as our perceptual systems have evolved to efficiently process the predictable redundancies of the scenes in which we evolved^11,23,46^. The Fourier slope is a straightforward way of measuring those redundancies and has been shown to correlate with aspects of visual processing. For instance, people are able to discriminate different textures based solely on differences in their Fourier slope^26^. Moreover, neurons active in the early stages of visual processing in vertebrates are specialized to respond to contrast at specific spatial frequencies. Notably, these specializations correspond to the spatial frequencies that occur with the highest energy in natural scenes^11,12,47,48^. Stimuli that most closely mimic the energy of spatial frequencies in natural scenes, which is quantified by the Fourier slope, should thus be most efficiently processed, as they generate a sparse neurological code that stimulates a small number of highly specialized neurons^10^.

Although it is now well supported that, in humans at least, efficient information processing is rewarded with pleasure, why this occurs is still unknown^18,49,50^. One explanation is a “processing bias” (i.e. Renoult & Mendelson)^5^, which supposes that this reward first evolved as an adaptation to inform the brain that information gathering is going smoothly, or that the environment is familiar^58^. Such a processing bias may secondarily be exploited by communication signals that are efficiently processed (e.g., that mimic the visual statistics of a familiar environment) and thereby elicit pleasure^5^. According to this hypothesis, male sexual displays in darters would have exploited female pleasure that originated as a general adaptation not associated with mate choice. Interestingly, we found a correlation between the Fourier slope of darter patterns and that of their habitat only in males, not in females, consistent with a hypothesis that male signals have evolved to be attractive, whereas female signals have not.

The difference we found between the sexes is puzzling, however, in the context of camouflage, which is the most obvious reason to expect a correspondence between the visual statistics of animal patterns and their environment. If mimicking the Fourier slope of the environment was driven by selection for camouflage, we would expect females, which are also subject to selection by predation, to follow the same pattern as males. Moreover, as males in breeding condition appear to contrast their environment, presumably maximizing detectability to mates or competitors, it seems unlikely that their visual displays are adapted for camouflage. Nevertheless, our analyses consider luminance contrast, rather than chromatic contrast; thus, the luminance patterning of males may be a camouflaged backdrop against which conspicuous colors are displayed. The attention that male colors draw from predators may add increased pressure to be otherwise camouflaged. Although the answer is not yet clear, the interplay of natural and sexual selection in driving the evolution of complex visual patterns remains an open question upon which the spatial statistics of animal patterns and their habitats could be fruitfully brought to bear.

Last, our finding that variation in the Fourier slope of darters is consistent with variation in species’ preferred habitat may have implications for human aesthetics. The Fourier slope has been found to vary between different categories of stimuli (e.g., buildings, natural landscapes, anthropogenic objects), and between terrestrial and aquatic environments (for a review, see Pouli et al.)^26^; however, the extent to which natural human habitats vary in Fourier slope has not yet been explicitly addressed. Different terrestrial biomes (e.g., tropical forest, desert, seashore) very likely have different Fourier slopes; thus, quantifying how they co-vary with works of art and regional aesthetic preferences may further contribute to understanding the mechanisms driving aspects of cultural evolution and diversification.

In conclusion, while our study does not directly investigate perceptual processing, it provides a plausible explanation for how sensory integration beyond photoreceptors can drive signal design through an environmentally mediated process. Through the lens of Fourier analysis, we have provided evidence that male visual signals correspond to the visual statistics of their habitats, suggesting that post-retinal visual processing is a plausible explanation for certain aspects of signal design. Our hypothesis is a novel extension of sensory drive, and our methods provide a new approach for testing the role of sensory drive in the evolution of visual patterns. Because the animal patterns we studied are likely used in mate attraction, our results also support key predictions of empirical aesthetics about the relationship between attractiveness and natural scenes. While empirical aesthetics was largely developed to explain human aesthetic preferences, we suggest that some of its principles extend beyond humans and provide a compelling hypothesis for how a complex trait can evolve in a predictable, environmentally dependent direction.

## Methods

### Darter Collection and Photography

We collected males and females of ten darter species from 23 sites distributed in Illinois, Kentucky, Maryland, Missouri, Pennsylvania, and Tennessee (*Etheostoma barrenense, E. blennioides, E. caeruleum, E. camurum, E. chlorosomum, E. gracile, E. olmstedi, E. pyrrhogaster, E. swaini, E. zonale*) (Supplementary Table 1). These ten species were chosen for inclusion based on their broad phylogenetic distribution and their preference for different classes of habitat: sand (*E. chlorosomum, E. olmstedi, E.pyrrhogaster*), boulder (*E. blennioides, E. camurum*), gravel (*E. caeruleum, E. zonale*), detritus (*E. gracile, E. swaini*) and bedrock (*E. barrenense*). At each site, we collected approximately 10 males and 10 females (males: 11.2 +/- 3.0 SD; females: 10.5 +/- 3.1 SD), which were subsequently photographed. Darters were caught by kick-seining and brought back to either the Hancock Biological Station in Murray, KY or University of Maryland Baltimore County. Fish were housed in aerated tanks and photographed within three days of capture. Immediately prior to photography, fish were euthanized in MS-222 and then fixed in 10% formalin with fins pinned erect for approximately 10 minutes. We then clipped the pectoral fin of each fish for an unobstructed image of their body pattern. Images were subsequently captured with a Canon EOS 5D Mark IV digital camera mounted to a Cognisys Stackshot system to ensure a fully focused image (see Supplementary Information for detailed photography methods).

### Habitat Photography

We collected images of habitats at sites where we captured darters using an Ikelite 200DL underwater housing for a Canon EOS 5D Mark IV digital camera. Each darter species was assigned to a habitat class (sand, gravel, boulder, detritus, or bedrock) based on where darters were observed, as well as on literature describing darter microhabitat preferences^52–54^. Each habitat class was represented by a minimum of 100 images representing a minimum of two sites. All images were collected in clear, shallow water on sunny days between 10:00 and 15:00, and when water turbidity was low (see supplementary information).

### Image Conversion to Darter Color Space

For all images, RAW files were converted to 16-bit tiff files using the rawpy python API. We converted RAW files to RGB triplets without any spatial interpolation, gamma correction or white-balance to maintain linearity. Images were then converted into a darter colorspace. The generation of the darter colorspace was done by first characterizing the sensitivity functions of our camera, and then using known darter visual sensitivities to generate a mapping from camera space to darter space^55^. The sensitivity functions for the camera were estimated using a monochrometer and a calibrated spectrometer, which ensured that each color channel was linear^56^. To model darter color vision, we generated a dichromatic model using cone sensitivities peaking at 525nm and 603nm (darters lack a short wavelength sensitive cone class). Since the cone sensitivities for all species in this study are not currently known, we used the same color vision model for all species. Variation between darter species in cone sensitivities is known to be relatively minor and unlikely to affect the outcome of our analysis^55^. We converted camera space to darter space by minimizing the difference between the camera model and the cone model, using a second order polynomial function of RGB values. We then converted all images (fish and habitat) from color space to luminance space by summing the two color channels. This pooling of color channels closely mimics how vertebrate brains are thought to extract luminance information^57^.

For darter images, we cropped out the region on the flank of each fish directly below the second dorsal fin. From the set of cropped images, we determined the largest square area that would fit in every cropped image, which was found to be 200 × 200 pixels. For each darter image, we then randomly sampled a region of this size from the flank of the fish. Habitat images were reduced from their original dimensions of 2251 × 3372 to 800 × 1200. We then randomly sampled each habitat image four times with a 400 × 400 pixel square. Since the size of the habitat images is greater than the size of the darter’s flank, using a larger box size reduces variability in lower frequency coefficients. Additionally, we tested the effects of various box sizes, and found our results robust to these changes (see supplemental information).

### Fourier Analysis

To compute the slope of the Fourier power spectrum for each image, we followed standard methods in empirical aesthetics^21,42,58^. We calculated the two-dimensional Fast Fourier Transform with a Kaiser-Bessel window using parameter α = 2 to minimize edge artifacts^26^. We then transformed the Fourier space to the power spectrum and estimated the radial average of the power spectrum. To eliminate edge effects and high frequency noise, we only included spatial frequencies between 10 and 110 cycles per image. Since the Fourier power spectrum has a greater spatial granularity at higher frequencies, we binned each power spectrum between 10 and 110 cycles for darter images and between 10 and 200 cycles for habitat images, with 20 bins for each. This ensures that our slopes were calculated to give equal weight across the frequency range. We then estimated the slope of the power spectrum using a linear regression on the bin values, using a custom Python script.

### Statistical Analysis

To ensure that images of species-typical habitat were representative of their class, we pooled images of each habitat class across multiple sites. We then compared the Fourier slope across habitat classes and across darter species (males and females analyzed separately) using ANOVAs. To examine the relationship between the Fourier slope of habitats and that of darter patterns, we used generalized linear mixed models. We computed this model using the R package MCMCglmm^59,60^. Our model predicted the value of the Fourier slope of each individual fish based on the slope of their habitat. Capture site and phylogeny were included as random effects. The phylogenetic tree of the ten studied species was inferred from a previously published molecular phylogeny (accessed via TreeBASE)^30,61^. We ran the model using the uninformative Inverse-Wishart prior for 1,000,000 iterations, with a 10,000 iteration burn in and 50 iteration thinning.

### Data Availability

All code used to compute the slope of the Fourier power spectrum can be accessed via github at https://github.com/svhulse/Fourier-Analysis. The computed slopes for every image used can also be accessed via github under the same repository. Any images used in this study are available upon request.

## Acknowledgements

We would like to thank Dr. Thomas Cronin for help with camera calibrations, Matthew Dugas, Natalie Roberts and Rickesh Patel for assistance with field collections, and the Hancock Biological Station for providing a home base for field work and photography. This work is supported by the Natural Science Foundation grant IOS-1708543.

## Author contributions

T.C.M. and J.P.R. conceived and designed this study. S.V.H. collected fish with assistance from T.C.M. and J.P.R., and was responsible for all photography. Additionally, S.V.H. wrote all python code and performed all analyses. All authors worked to write and edit the manuscript.

## Materials & Correspondence

For any additional information, data, or images used in this study, please contact S.V.H.

