## Supplementary material for "Sexual signals of fish species mimic the spatial statistics of their habitat: evidence for processing bias in animal signal evolution": Supplimentary Information

**Supplementary Information**

**Methods**

**Supplemental Methods**

**Permits**

Permits for scientific collection were obtained through the Kentucky Department of Fish and Wildlife Resources (Permits SC1711119 and SC1811151), Tennessee Wildlife Resources Agency (Permits 1052 and 1424), Illinois Department of Natural Resources (Permits A17.6089 and A18.6089), Missouri Department of Natural Resources, Louisiana Department of Natural Resources (Permit SCP 183), Mississippi Department of Wildlife, Fisheries and Parks (Permit 0219184) and Maryland Department of Natural Resources (Permit # SCP201747).

**Image Capture**

For habitat images, each image was captured by holding our camera near to the bottom of the stream, and taking pictures facing multiple directions. This was repeated across each field site until the area where we found darters was photographed to the complete extend that conditions allowed, and until each habitat has at least 100 photographs. We also took images of a waterproof white balance card (DGK Color Tools WDKK Waterproof Color Chart) at each site. We photographed all darters within the three days following capture. Each fish was first euthanized with MS-222, and then pinned with fins erected in a bath of 10% formalin. After approximately 10 minutes, fish were removed from the formalin and placed in an open-topped cylindrical glass photography arena with water at a depth of 2cm. The arena was surrounded by diffuser paper and illuminated by three Canon 270EX II flashes spaced equidistant around the arena. Each fish was placed in the arena with the camera facing perpendicular to their side. We then photographed each fish using a Canon EOS 5D Mark IV digital camera with a Canon EF 100mm f/2.8L macro lens attached. The camera was mounted on a Cognisys Stackshot Extended Macro Rail to enable automated focus stacking. Prior to imaging, the camera’s position on the rail was set so that the most proximate part of the darter was in focus. The macro rail was then set to capture an image every 0.5mm, until the most distal part of the fish was in focus. During every photography session, we also captured an image of a white balance card in the photography arena.

**Image Processing**

To retain the high dynamic range and linearity of the RAW image files, we used a custom python script using the library libraw to extract image data from each Canon RAW file. We applied no gamma correction, or white balance. Additionally, each image was reduced to half size to avoid nonlinearities associated with demosaicing algorithms. For darter images, we then combined each image stack to a single image using Zerene Stacker with the DMap algorithm.

**Tables**

Table 1: Locations for all field sites where darters were collected, number of individuals collected for each species, as well as their habitat classifications, and the references used to determine the habitat classification.

| Species | Sample Sites | GPS Coordinates (E, N) | Habitat | Number Collected (females, males) | References |
| --- | --- | --- | --- | --- | --- |
| *E. barrenense* | EF Barren River  Line Creek  Trammel Creek | 36.7459. -85.6967  38.6069, -85.7458  36.7396, -87.2896 | Bedrock | 18, 12  12, 18  13, 12 | Etnier and Starnes, 1993  Kuehne and Barbour, 2014 |
| *E. blennioides* | MF Red River  Jordan Creek  Boone Creek | 37.7815, -83.6824  40.3533, -87.5502  38.2582, -91.2832 | Boulder | 12, 8  7, 11  12, 11 | Etnier and Starnes, 1993  Kuehne and Barbour, 2014 |
| *E. caeruleum* | MF Red River  Trammel Fork  Salt Fork | 37.8149, -83.7187  36.7520, -86.2872  40.0829, -87.7806 | Gravel | 12, 15  12, 12  11, 11 | Etnier and Starnes, 1993  Kuehne and Barbour, 2014 |
| *E. camurum* | SF Kentucky River  MF Kentucky River  MF Vermillion River | 37.3381, -83.6880  37.0776, -83.3926 | Boulder | 12, 11  10. 11  9, 10 | Etnier and Starnes, 1993  Kuehne and Barbour, 2014 |
| *E. chlorosomum* | Old Town Creek | 36.3082, -88.4488 | Sand | 9, 12 | Etnier and Starnes, 1993  Kuehne and Barbour, 2014 |
| *E. gracile* | Embarras River  Skillet Fork  Brush Creek | 38.9074, -87.9078  38.7088, -88.6645  38.5350, -88.6121 | Detritus | 10. 10  5, 11  4, 12 | Petersons, 1991  Kuehne and Barbour, 2014 |
| *E. olmstedi* | M Patuxent River  Rock Creek | 39.1680, -76.8833  39.1510, -77.1036 | Sand | 11, 11  11, 9 | Etnier and Starnes, 1993  Kuehne and Barbour, 2014 |
| *E. pyrrhogaster* | Barnes Fork  Old Town Creek | 36.2430, -88.2874  36.3082, -88.4488 | Sand | 12, 12  11, 11 | Bailey and Etnier, 1988  Etnier and Starnes, 1993  Kuehne and Barbour, 2014 |
| *E. swaini* | Scarborough’s Creek  Moaks Creek  Myers Creek | 30.9379, -89.7782  31.372, -90.4395  31.4337, -90.42 | Detritus | 11, 12  11, 12  5, 11 | Etnier and Starnes, 1993  Kuehne and Barbour, 2014 |
| *E. zonale* | Line Creek  MF Kentucky River  Little Sugar Creek | 36.6519. -85.8204  37.0776, -83.3926  41.5224, -80.0498 | Gravel | 16, 11  12, 10  7, 15 | Etnier and Starnes, 1993  Kuehne and Barbour, 2014 |

Table 2: Average Fourier slopes for males and females of each species as well as the significance of the intersexual difference (Bonferroni correction)

| Species | Mean (male) | Mean (female) | Corrected p-value | t-value | df |
| --- | --- | --- | --- | --- | --- |
| *E. barrenense* | -3.052 | -3.329 | 1.124e-5 | 5.275 | 79.23 |
| *E. blennioides* | -3.173 | -3.357 | 0.7136 | 1.836 | 58.79 |
| *E. caeruleum* | -3.419 | -3.375 | 1 | -0.6934 | 70.17 |
| *E. camurum* | -3.175 | -3.183 | 1 | 0.1109 | 59.99 |
| *E. chlorosomum* | -2.643 | -2.689 | 1 | 0.4725 | 15.56 |
| *E. gracile* | -3.129 | -3.121 | 1 | -0.1057 | 34.56 |
| *E. olmstedi* | -3.029 | -3.389 | 4.041e-5 | 5.361 | 38.79 |
| *E. pyrrhogaster* | -2.645 | -3.195 | 1.584e-6 | 6.266 | 42.41 |
| *E. swaini* | -3.086 | -3.306 | 0.002144 | 3.947 | 58.55 |
| *E. zonale* | -3.056 | -3.29 | 0.02655 | 3.123 | 65.73 |
